# Neuronal expression of herpes simplex virus type-1 VP16 protein induces pseudorabies virus escape from silencing and reactivation by activating Jun

**DOI:** 10.1101/2023.01.13.524029

**Authors:** Zhi-Shan Hsu, Esteban A. Engel, Lynn W. Enquist, Orkide O. Koyuncu

**Affiliations:** Department of Molecular Biology, Princeton University, Princeton, NJ; Princeton Neuroscience Institute, Princeton University, Princeton, NJ; Microbiology and Molecular Genetics Department, University of California Irvine, Irvine, CA; Spark Therapeutics, Philadelphia, PA

## Abstract

Alpha herpesvirus (α-HV) particles enter their hosts from mucosal surfaces and efficiently maintain fast transport in peripheral nervous system (PNS) axons to establish infections in the peripheral ganglia. The path from axons to distant neuronal nuclei is challenging to dissect due to the difficulty of monitoring early events in a dispersed neuron culture model. We have established well-controlled, reproducible, and reactivateable latent infections in compartmented rodent neurons by infecting physically isolated axons with a small number of viral particles. This system not only recapitulates the physiological infection route, but also facilitates independent treatment of isolated cell bodies or axons. Consequently, this system enables study not only of the stimuli that promote reactivation, but also the factors that regulate the initial switch from productive to latent infection. Adeno associated virus (AAV) mediated expression of herpes simplex type 1 (HSV-1) VP16 alone in neuronal cell bodies enabled the escape from silencing of incoming pseudorabies virus (PRV) genomes. Furthermore, expression of HSV VP16 alone reactivated a latent PRV infection in this system. Surprisingly, expression of PRV VP16 protein supported neither PRV escape from silencing nor reactivation. We compared transcription transactivation activity of both VP16 proteins in primary neurons by RNA sequencing and found that these homolog viral proteins produce different gene expression profiles. AAV transduced HSV VP16 specifically induced expression of proto-oncogenes including Jun and Pim2. In addition, HSV VP16 induces phosphorylation of Jun in neurons, and when this activity is inhibited, escape of PRV silencing is dramatically reduced.

## Introduction

Alpha herpesviruses (α-HV) are common pathogens of mammals. Herpes simplex type 1 (HSV-1 and herpes simplex type 2 (HSV-2) infect the vast majority of the adult human population. These viruses are the causative agents of cold sores, genital herpes, herpes stromal keratitis, and encephalitis [1-3]. All α-HV infections begin as a productive infection in the epithelial cells of mucosal surfaces (e.g., the nasal-pharyngeal cavity, genitals) [2, 4, 5]. Some of the progeny virus particles invade the innervating axons of PNS neurons. Viral particles then engage microtubule-based molecular motors in these long axons to reach neuronal nuclei [6-9]. Viral genomes reaching PNS nuclei in the peripheral ganglia usually are silenced and remain quiescent for long periods of time, a state often referred to as latency.

PNS neurons have specialized signaling and gene expression patterns that utilize the highly polarized morphology for optimal function. Axonal biology and efficiency of long-distance transport of viral particles influence whether the infection of α-HVs in neurons will be productive or latent. The sequence of events starting with axonal invasion and ending with the genome in PNS neuronal nucleic is challenging to dissect due to the difficulty of isolating early events in axons independent from later evens in neuronal nuclei. Data from animal models and cultured primary neuron models revealed that the decision to replicate or enter latency depends on the presence of viral outer tegument proteins, particularly VP16 (virion protein 16, aka UL48) protein [7, 10-12]. Incoming VP16 proteins delivered by virus particles interact with host transcription factors in the nucleus to activate immediate early (IE) viral gene expression to initiate productive infection [12-14]. However, VP16 is an outer tegument protein that is not co-transported with capsids to the neuronal nucleus. It is still not well known whether VP16 in the tegument undergoes retrograde transport in axons alone or with other outer tegument proteins [15-19]. Such separate transport of outer tegument and viral capsids in axons may lead to their asynchronous arrival at the neuronal nucleus and could bias the infection mode toward latency [7, 20]. The VP16 gene is classified as a late gene, being efficiently transcribed after DNA replication begins. However, recent studies provided evidence that regulatory features in the 5`UTR of the VP16 gene, close to the promoter region ensures pre-immediate early (pre-IE) expression of VP16 [11, 21]. The expression and transactivation activity of VP16 is essential to initiate the gene expression cascade transitioning from an established latent infection to a productive state (i.e., reactivation) [11, 22, 23]

In this paper, we investigated the effect of the related VP16 proteins encoded by PRV or HSV-1, on PRV escape from genome silencing. PRV is a varicellovirus from the *alphaherpesvirinae* subfamily and is a well-known veterinary pathogen [24]. PRV and HSV-1 share common strategies to infect and invade the nervous system. VP16 is one of the most abundant proteins (app. 1000-1500 copies) in both HSV-1 and PRV tegument [16, 25]. The functions of the HSV-1 VP16 protein are well known, but PRV VP16 is less well studied. While both proteins are required for transcription of immediate early viral genes, how the proteins affect neuronal gene transcription remains unclear. There are some significant differences between the structure and function of the homolog viral proteins. The HSV-1 VP16 protein has 490 amino acid residues, whereas PRV homolog contains 413. There is no significant sequence similarity, but some motifs are conserved such as the serine residue (S375) critical for reactivation of HSV-1 (Fig 1A and B). PRV and HSV-1 also differ in the number of immediate early (IE) genes that are encoded: HSV-1 has 5, whereas in PRV has only 1. PRV IE180 is the only IE gene required for DNA replication and RNA transcription.

**Figure 1.**
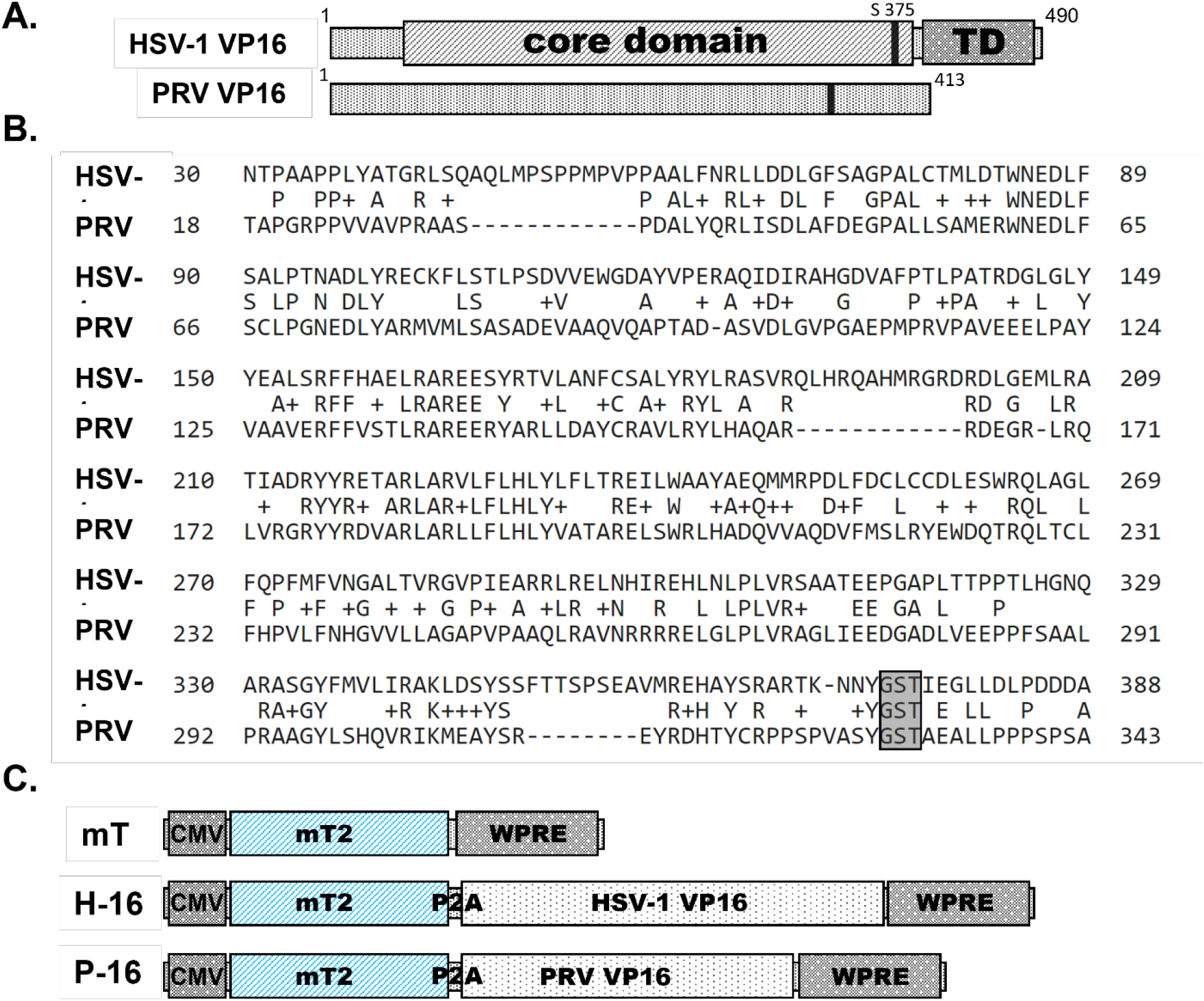
Comparison of VP16 protein coding sequences of HSV-1 and PRV. (A) Diagrams of HSV-1 and PRV VP16 coding sequences (core domain and transactivation domain-TAD-are shown for HSV-1 VP16). (B) Comparative analysis of the protein sequences of HSV-1 VP16 (Query) and PRV VP16 (Subject) using NCBI BLAST. The boxed section contains the Ser375 residue of HSV-1 VP16 and the analogous Ser330 residue in PRV VP16. (C) Constructs designed for adeno associated vector (AAV) expression. mT: mTurqoise2, cmv: cytomegalovirus promoter, P2A: self-cleaving 2A sequence, WPRE: Woodchuck Hepatitis Virus Posttranscriptional Regulatory Element

To establish a well-controlled, reproducible, and reactivatable α-HV latent infection in cultured PNS neurons without inhibiting DNA replication, we cultured rat superior cervical ganglionic (SCG) neurons in modified Campenot chambers with three compartments (tri-chambers), and infected isolated axons with a small number of viral particles. Tri-chambers physically and fluidically separate neuronal cell bodies from axons during the establishment of neuronal polarity and maturation. The advantage of this approach is that it not only facilitates analysis of the physiological infection route (from axons to cell bodies), but also it enables treatment of isolated cell bodies or axons separately to activate or inhibit target pathways. This model further enables study of the factors that regulate the initial switch from productive to latent infection. Using this model, previously we investigated viral and cellular factors regulating productive versus latent PRV infection [8]. Activation of neuronal protein kinase A (PKA) and cJun-N-terminal kinase (JNK) in the cell bodies led to a slow escape from silencing. Interestingly, when viral tegument proteins were delivered to cell bodies either by capsid-less light particles or UV inactivated virus particles, the axonal infection rapidly switched to a productive mode independent of cellular PKA and JNK pathways [8].

Here we used adeno-associated virus (AAV) vectors to deliver the individual proteins (either HSV-1 or PRV VP16) to compartmented SCG neurons before initiating a latent PRV infection from axons. Surprisingly, HSV-1 VP16 alone potently induced a rapid escape from PRV genome silencing. On the other hand, PRV VP16 alone was not able to induce such an escape. Moreover, AAV delivery of HSV-1 VP16, but not PRV VP16, to neurons after PRV latency was established, reactivated productive PRV infection. We further compared the transactivation activity of neuronal genes of both viral proteins by transducing primary neurons with the AAV vectors. We found that these homolog viral proteins induced transcription of different subsets of neuronal genes. HSV-1 VP16 specifically activated transcription of proto-oncogenes, particularly Jun, while PRV VP16 increased levels of dual specificity phosphatase 4 (Dusp4; MKP-2) and Adenylate cyclase activating polypeptide 1 (Adcyap-1; PACAP) transcripts. We confirmed that HSV VP16 induction of PRV escape from silencing is dependent on the Jun pathway. These comparative experiments may offer new insight into the molecular biology of alpha herpesvirus latency and reactivation and how VP16’s role in these processes affects the expression of cellular genes.

## Results

### Construction of recombinant AAV vectors expressing HSV-1 VP16 (H-16) or PRV VP16 (P-16)

To achieve high transduction efficiency of VP16 into neurons, we constructed adeno associated virus vectors (AAV) expressing either HSV-1 or PRV VP16. We chose AAV serotype PHP.eB that has shown broad tropism in primary rodent neurons. Recombinant AAV vectors are non-toxic, rarely integrate into the host genome, and expression of the transgene increases over time giving ample time to monitor transgene expression [26, 27]. For monitoring transduction and expression efficiencies, we cloned the mTurqoise2 gene upstream of the VP16 sequence separated by the p2A cleavage sequence. We did this to avoid fusing the reporter to VP16 sequences with potential interference of the fluorescent reporter with the transactivation domain, or any other protein interaction domain in VP16. AAV plasmids coding for mTurqoise2 only (mT), mTurqoise2-P2A-HVP16 (H-16), and mTurqoise2-P2A-PVP16 (P-16) were synthesized by Genscript and used to produce AAV-PHP.eB vectors (see Materials and Methods) (Fig 1C). Accumulation of mTurqoise2 protein started 2 days after the mT transduction, whereas detectable fluorescent protein accumulated in neuronal cell bodies only after 3 days after H-16 and P-16 transduction (Fig 2A). To assess the cleavage efficiency of the protein fragments, we harvested SCGs at 5 days post transduction (dpt) and analyzed proteins by immunoblotting. A rabbit polyclonal antibody detecting PRV VP16 protein was made by Genscript. This antibody detected a 50 kDa band in the P-16 transduction sample (Fig 2B). Anti-HSV-1 VP16 antibody (Abcam) detected an approximately 55 kDa band in the H-16 sample. Neither of the antibodies cross-reacted with the homolog viral proteins. mTurqoise2 was detected by an anti-GFP antibody in all three samples. An uncleaved fusion protein was detected approximately at 75 and 80 kDa, in P-16 and H-16 transduction samples respectively. The intensity of these bands was lower than the VP16 and mTurqoise2 bands indicating that most of the fusion protein is cleaved at this time point.

**Figure 2.**
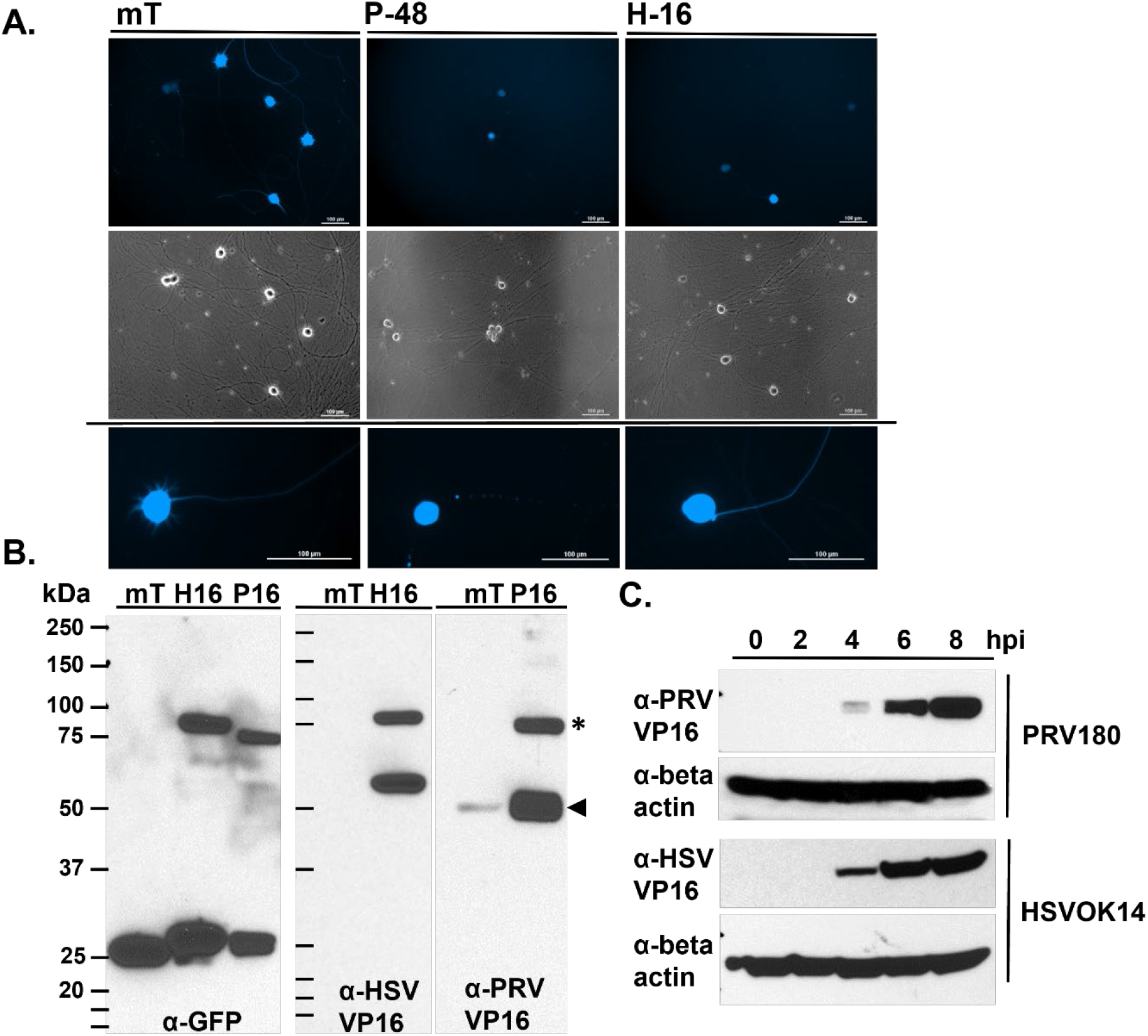
Expression analysis of VP16 from recombinant AAV vectors and after viral infection. (A) mTurqoise2 expression levels of AAV-transduced SCGs after 96 hours. Neurons, from left to right, were transduced with AAVs encoding mTurqoise2 (mT), PRV VP16 (P-16), and HSV VP16 (H-16). Scale bars represent 100 μm. (B) Western Blot analysis of neurons transduced with the AAV vectors as in (A). Cells were lysed and protein expression levels were investigated for HSV and PRV-VP16 using corresponding antibodies. The expected sizes of HSV VP16 is 56 kDa and PRV VP16 is 50 kDa (arrowhead). Uncut protein (mT-P2A-VP16) shown with an asterix. (C) Western Blot analysis of Rat2 cells infected with HSV-1 OK14 or PRV180 at MOI of 10. Cells harvested at a different time-points after infection (hpi: hours post infection) as indicated. Endogenous VP-16 expression was monitored during virus infection. Beta-actin is used as a loading control.

We also tested the capacity of antibodies to recognize VP16 proteins expressed after virus infection (Fig 2C). Rat2 cells either were mock infected or infected with HSV-1 OK14 or PRV180 recombinants expressing mRFP-VP26 at an MOI of 10. Infected cells were harvested at 2, 4, 6 and 8 hours post infection (hpi). Both VP16 proteins expressed either during HSV-1 or PRV infection showed comparable steady state levels, starting 4 hpi and accumulating over time (Fig 2C). Immunofluorescence analysis also showed punctate VP16 staining at 2 hpi, mostly overlapping with the capsid signal (mRFP-VP26) in both HSV-1 and PRV infection (Fig 3). As expected, both PRV- and HSV-infected Rat2 cells exhibited nuclear localization of VP16 at 4 hpi. At each subsequent time point, more VP16 was seen in the cytoplasm and, at 8 hpi, most of the tegument protein was cytoplasmic. From these data, it appears that VP16 proteins of HSV-1 and PRV are expressed in a similar time frame with a comparable subcellular localization in infected cells.

**Figure 3.**
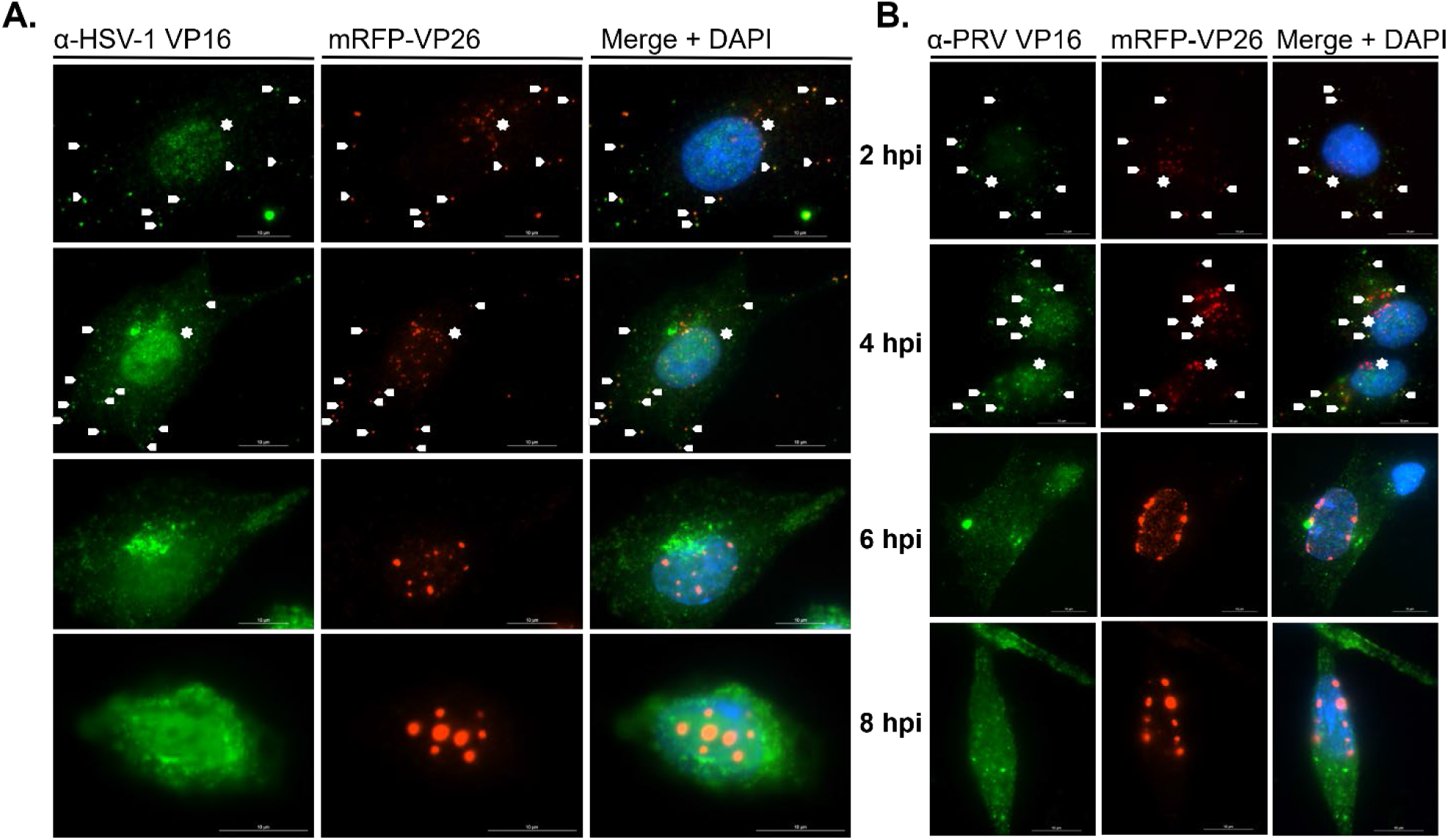
Subcellular localization of VP16 during HSV-1 OK14 (A) or PRV 180 (B) infection in Rat2 cells. Infected cells were fixed every 2 hours and were stained with corresponding VP16 antibodies. VP26 is expressed as mRFP fusion in these recombinant viruses. Hpi stands for hours post infection. Scale bars represent 10 μm. Arrows show intact (dual color) virus particles whereas stars point to green only particles most probably light particles at 2 and 4 hpi.

### HSV-1 VP16 but not PRV VP16 expression in SCGs enables PRV fast escape from silencing

To investigate whether HSV-1 or PRV VP16 protein alone is capable of inducing PRV escape from silencing, we performed a `complementation assay` in compartmented neuronal cultures as we previously described [8]. Briefly, we determined the infectious PRV dose infecting axons that results in latent infections in the neuronal cell bodies in the distant S-compartments. Before infecting axons with PRV, we transduced neurons in the S-compartments with AAV vectors mT, H-16 or P-16. We confirmed transgene expression by monitoring mTurqoise2 accumulation. Four days after AAV transfection, we infected isolated axons in the N-compartments with PRV180 at an MOI of 0.01 and monitored mRFP-VP26 accumulation in the cell bodies over time. As controls, we treated cell bodies in the S-compartments with (UV inactivated PRV (UVPRV) or PRV light particles (LP), while infecting axons with PRV180, as both conditions result in fast escape from genome silencing [8]. mRFP-VP26 accumulated in cell bodies as early as 48 hpi in chambers transduced with H-16. By 3 dpi, all the H-16 transduced chambers had productive PRV infection that spread all among neuronal cell bodies in S-compartments. At this time point, control dishes treated with UVPRV and LP also had productive infection as deduced by mRFP capsid accumulation and spread among cell bodies. We saw no evidence of productive infection in the chambers transduced with P-16 and mT even at 5 dpi.

### HSV-1 VP16 expression in SCGs reactivates latent PRV infection

Latent PRV infections were established in cultured neurons by infecting isolated axons at MOI of 0.01 with PRV180. Individual chambers were screened every 3 days for mRFP-VP26 expression and accumulation reporting the late phase of productive infection. We did not detect any mRFP-VP26 signal in these chambers for 2 weeks indicating establishment of *in vitro* latency. Reactivation assays were performed by transducing cell bodies in the S-compartments with mT, H-16 and P-16. mRFP-VP26 accumulation in the cell bodies was monitored over time (Fig 5A). In all the chambers transduced with H-16, mRFP-VP26 signal was detected as early as 2 dpt, and reactivated virus infections spread among the S-compartment cell bodies by 4 to 6 days (Fig 5B). PRV reactivation was not detected in any of the mT or P-16 transduced chambers (Fig 5C). These chambers were imaged for mRFP-VP26 for 10 days.

**Figure 4.**
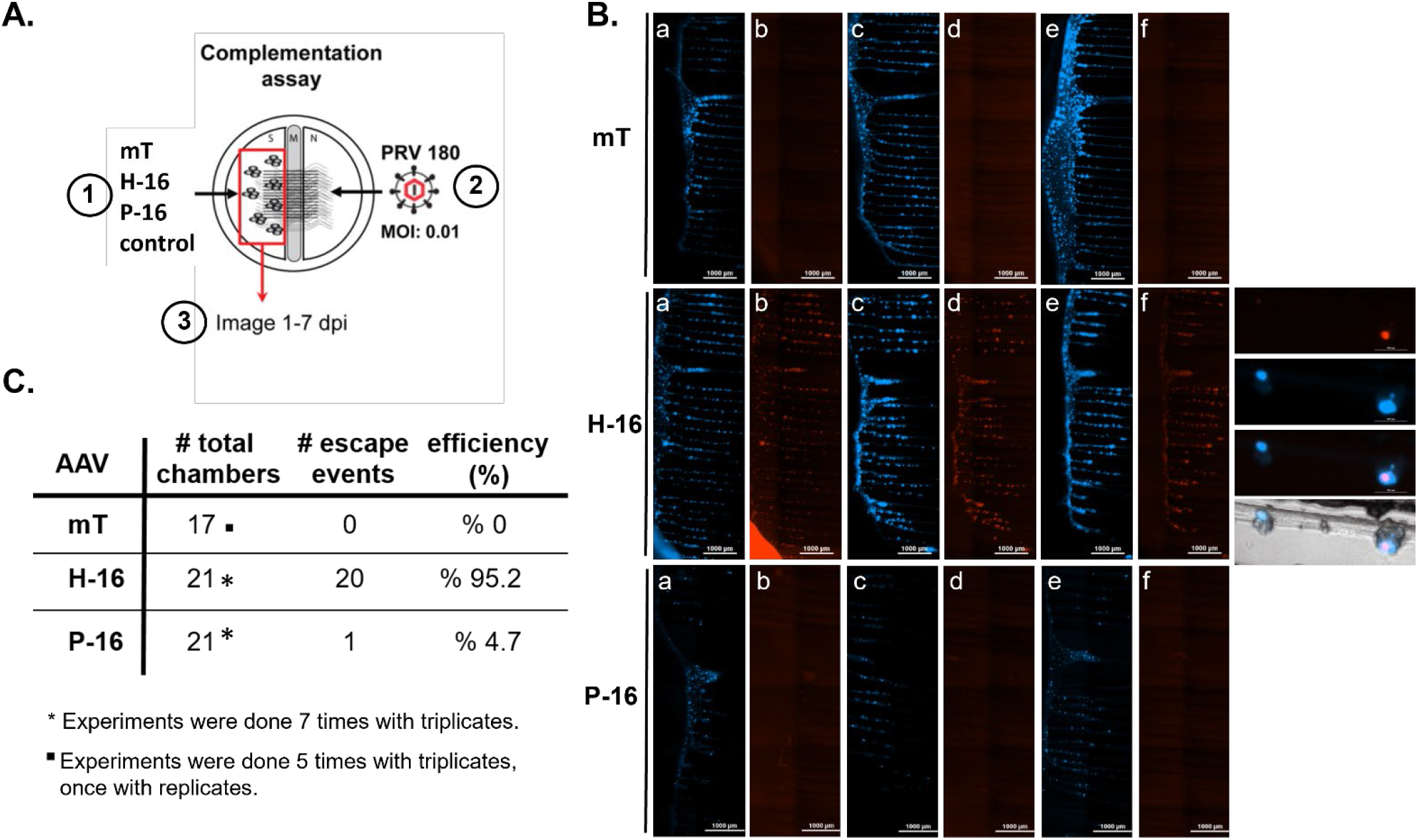
PRV Escape from silencing experiment. (A) Illustration of the complementation assay: 1) S-compartments were transduced with AAV vectors expressing mT, H-16, P-16 or kept as control 2) 3-days post transduction, N compartments were infected with PRV180 at an MOI of 0.01. 3) S-compartments were imaged starting from 24 hpi for the next 7 days. (B) Whole-S-compartment tiled images are shown for mT and VP26-mRFP expression at 3 dpi (H-16 insets shows detection of red fluorescent capsids proteins at 2 dpi) (C) Table summarizing all the experimental conditions and results of the experiment.

**Figure 5.**
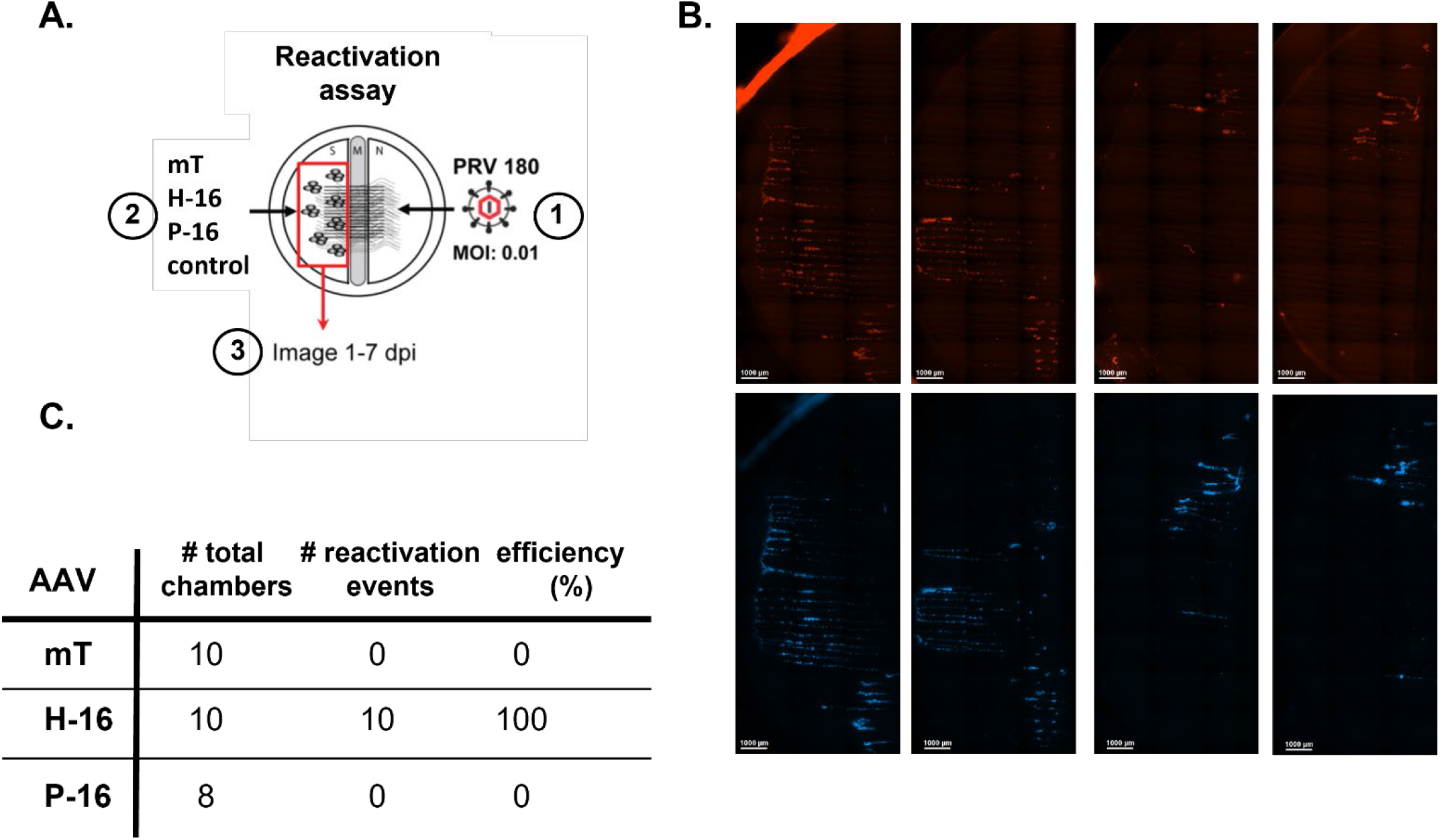
PRV Reactivation experiment. (A) Illustration of the complementation assay: 1) N compartments were infected with PRV180 at an MOI of 0.01 2) 10 dpi, S-compartments were transduced with AAV vectors expressing mT, H-16, P-16 or kept as control 3) S-compartments were imaged starting from 24 hpi for the next 7 days. (B) Whole-S-compartment tiled images are shown for VP26-mRFP expression at 4 dpi (C) Table summarizing all the experimental conditions and results of the experiment.

### Comparing the neuronal gene transactivating activity of HSV-1 VP16 to PRV VP16 in primary neurons using RNA seq

The effects of HSV VP16 and PRV VP16 proteins on neuronal gene expression have not been well characterized. RNA sequencing (RNA-seq) was performed on superior cervical ganglia (SCG) neurons transduced with one of three AAV vectors: mT, H-16 and P-16. Rat SCG neurons were cultured in 6-well dishes and transduced with the AAV vectors for 3 days. While an incubation time of 5 days would have ensured higher expression levels, we harvested transduced neurons at 72 hpt for RNA extraction to reduce the risk of triggering cellular stress responses due to the viral protein expression. 4 to 6 wells of a 6well dish were pooled together per sample for RNA extraction and experiment was repeated 3 times.

In the HSV VP16-transduced data, Jun and Pim2 were the most and second most enriched genes respectively (Fig 6A). The increase in the transcription levels of the proto-oncogene Jun was not surprising. Previous research has demonstrated that activation of the c-Jun N-terminal kinase (JNK) pathway increased HSV-1 reactivation and replication [28, 29]. Pim2 is a proto-oncogene that encodes kinases implicated in promoting cell survival and inhibiting apoptosis in multiple types of cancer.

**Figure 6.**
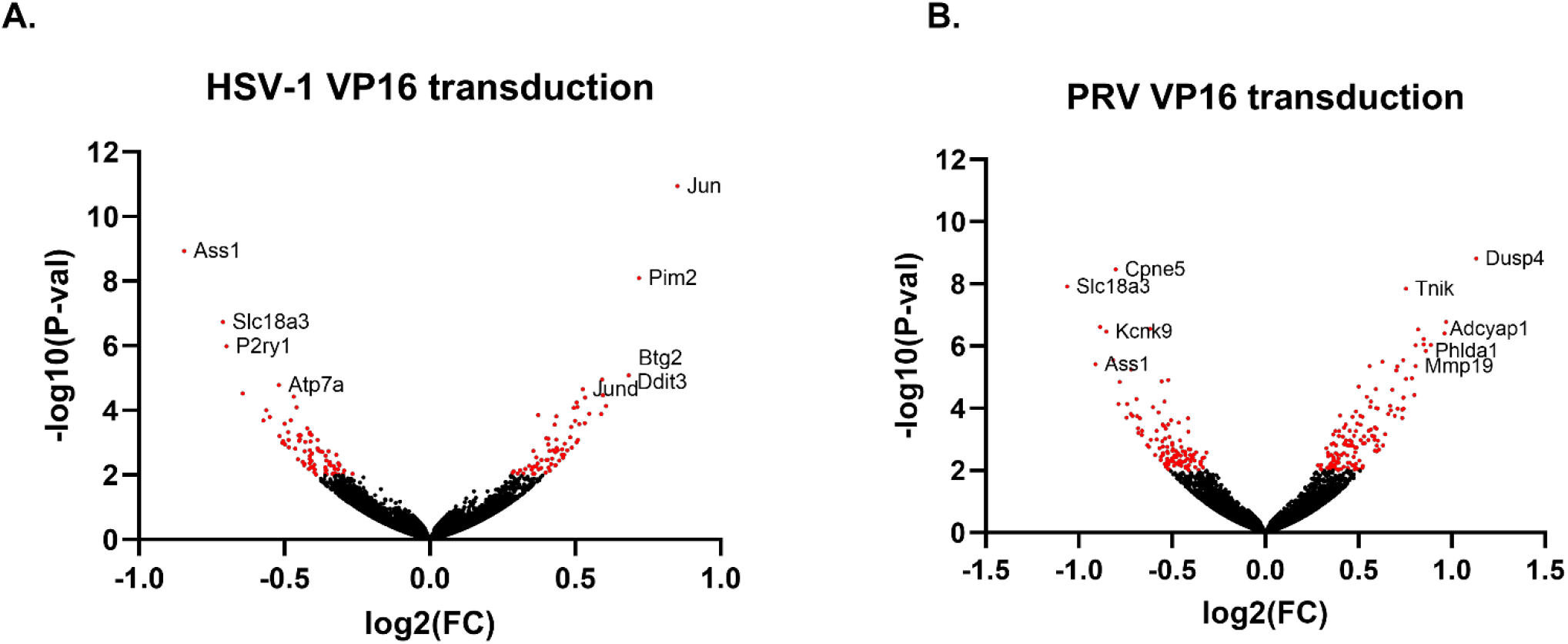
Volcano Plot for differential gene expression upon HSV-1 (A) or PRV (B) VP16 transduction. Neurons were transduced with HSV-1 or PRV VP16 or mTurqoise2 (mT) expressing AAVs for 3 days. Scattered points represent genes, the x-axis is the log 2-fold change VP16 vs mT expressing neurons, whereas the y-axis is the statistical significance in its differential expression. Red dots highlight genes significantly over or under -expressed after VP16 expression.

Dusp4 and Adcyap1 were the most expressed neuronal genes in the PRV VP16-transduced data. Dusp4 (dual specificity phosphatase 4) was identified as an inhibitor of the MAPK signaling pathway and was implicated in a range of roles from regulating muscle cell differentiation to controlling circadian rhythm. Adcyap1 (adenylate cyclase activating polypeptide 1), as the name implies, increases production of the second messenger cAMP by activating adenylate cyclase. Crem (cAMP response element modulator) was one of the few transcripts that was significantly expressed in both the PRV VP16- and HSV VP16-transduced samples.

### HSV-1 VP16-induced-PRV escape from silencing is mediated by activated Jun

Since we observed rapid escape from PRV genome silencing in H-16 transduced neurons, and Jun transcripts showed the highest significant increase after H-16 transduction, we determined if H-16 induced escape is dependent on activated Jun. First, we confirmed the activation of cJun after transduction of SCGs with H-16 in comparison to P-16 and mT transduction. We detected specific increased expression and phosphorylation of cJun only after H16 transduction (Fig 7A). Next, we transduced neurons in the S compartments with H-16, then 3 dpt, we treated the neuronal cell bodies in the S compartments with c-Jun N-terminal kinase inhibitors: JNKII (20 μM), JNK8 (10 μM), AS601245 (20 μM) or DMSO. 3 hours post treatment, we infected the axons in the N compartments with PRV180 at an MOI of 0.01. All of the JNK inhibitors severely reduced the efficiency of HSV-1 VP16-mediated escape from silencing of PRV genomes, indicating that HSV-1 VP16 protein induces rapid transcription of PRV genomes by activating the cJun signaling pathway that was not induced by PRV VP16 protein (Fig 7A, C).

**Figure 7.**
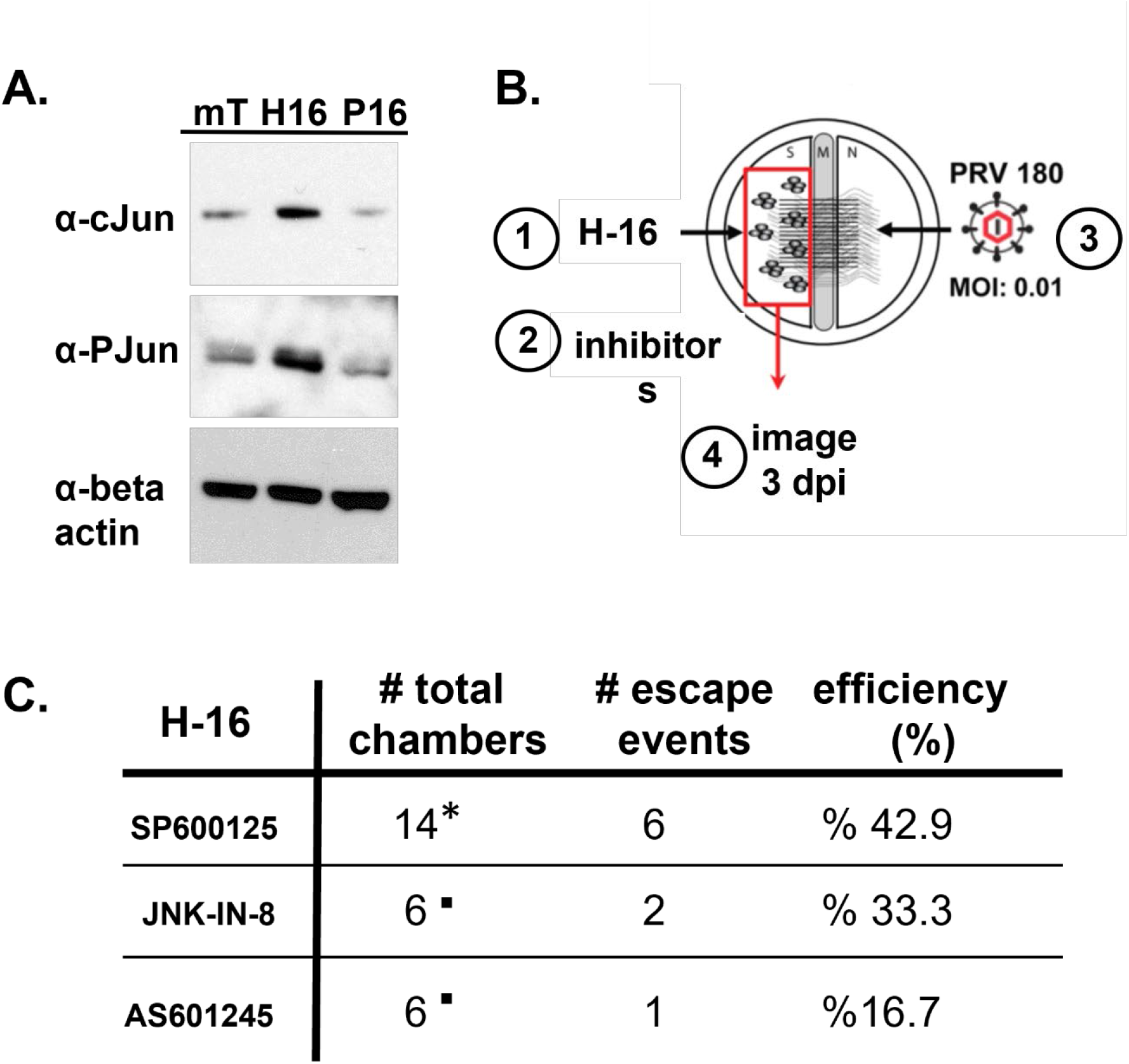
Escape from silencing experiment in the presence of JNK inhibitors. (A) Neurons were transduced with the AAV vectors expressing mT, H-16, P-16 for 4 days, and cJun and P-cJun protein levels were analyzed by western blotting. Beta-actin was used as loading control (B) Illustration of the complementation/inhibition assay: 1) S-compartments were transduced with AAV vector expressing HSV-VP16 2) 3-days post transduction, inhibitors were added in the S-compartments: SP600125 (10 μM), JNK-IN-8 (5 μM), AS601245 (20 μM) 3) Simultaneously, N compartments were infected with PRV180 at an MOI of 0.01 4) S-compartments were imaged starting from 24 hpi for the next 7 days. (C) Table summarizing all the experimental conditions and results of the experiment.

## Discussion

During latency, α-HV genomes are silenced yet retain the capacity to reactivate periodically to produce infectious viral progeny. The cue for reactivation is usually stress such as physical trauma, sunburn or fever [23, 30, 31]. Importantly, the threshold for reactivation is high such that it does not occur very often, Currently, host and viral factors that maintain latency and induce reactivation are not completely understood. One of the reasons is that the tissue architecture of PNS innervation is difficult to recapitulate in culture, which challenges dissecting the molecular mechanisms of reactivating cues.

Recently, we have devised a method to produce a reactivateable α-HV latent infection in vitro by using compartmented primary neurons [8, 32]. The compartmented neuronal culture system is well-controlled and recapitulates the natural route of latent infection (axon to cell bodies) eliminating the use of DNA synthesis inhibitors such as acyclovir. This model allows studying not only the reactivation but also the decision-making step early in the establishment of latency. Since the number of virus particles and conditions required to establish reproducible latency is determined, viral and cellular proteins and pathways that interfere with the establishment of a latent infection can be studied. By using this model, we found that activating neuronal protein kinase A (PKA) and cJun-N-terminal kinase (JNK) in the cell bodies induce a slow escape from silencing [8]. On the other hand, when viral tegument proteins were delivered to cell bodies, the infection rapidly switched to a productive mode independent of cellular PKA and JNK pathways [8]. In this paper, we investigated the effect of the tegument protein VP16 from two alpha herpesviruses, PRV and HSV-1, on the escape from silencing of incoming PRV genomes.

Live-cell imaging experiments examining the retrograde transport of monomeric red-fluorescent protein-tagged PRV capsids and green-fluorescent protein (GFP)-tagged tegument proteins in sensory neurons revealed that outer tegument proteins, such as UL47 (VP13/14), UL48 (VP16), and UL49 (VP22), were lost during viral entry and did not move along with capsids to the nucleus [6, 15, 33]. We and others hypothesized that such loss of the major transcriptional activator of viral genes during the retrograde transport might be the reason for silenced infection in PNS neurons [7, 30, 32].

Upon entry of the HSV genome into the nucleus, histones are loaded on the viral genome, and initial viral gene transcription depends on cellular epigenetic regulation. VP16 interacts directly with transcriptional activators and indirectly with epigenetic factors through host cell factor-1 (HCF-1) [34]. VP16 has a core region and C-terminal transcriptional activation domain (TAD). The core region is sufficient for VP16-induced transactivator complex formation, and the TAD is known to be critical for activation of transcription [12]. This transactivation is induced by interactions with multiple transcription factors including TATA-binding protein (TBP), transcription factor IIA (TFIIA), TFIIB, TFIID, and TFIIH [35-37]. In addition, the VP16 TAD interacts with subunits of Mediator [38, 39], implying a direct role in the regulation of the RNA polymerase II machinery.

Furthermore, studies using a series of virus mutants demonstrated that the transition out of the latent program (i.e., reactivation) is dependent on VP16 transactivation function [11, 40]. Collectively these data point that VP16 protein functions as a potent transactivator of viral IE genes in the sensory neuron, and the regulation of its transport to the neuronal cell body or its expression through the promoter regions determines the latency to productive infection transition [7, 10, 12, 14, 40-44].

Unlike HSV VP16, little is known about the host factors associated with PRV VP16 in neurons during infection, and whether the presence of VP16 protein in the cell bodies during initial infection is sufficient to start productive infection. Earlier reports confirmed that a PRV VP16 null mutant has a severe reduction in virus yield (almost 1000-fold) and shows defects in virion assembly in non-neuronal cells ([45, 46]. It was also noted that the VP16 protein in the virion can activate transcription of the sole PRV immediate early protein, IE180 [45]. Therefore, we hypothesized that PRV VP16, when present in the neuronal cell bodies, interacts with diverse neuronal proteins to activate viral gene expression, and induce the PRV productive cycle in our compartmented neuronal model of escape from genome silencing. We included the well-known HSV-1 VP16 in these assays to compare the transactivation potential of the two homolog alpha herpesvirus proteins. To achieve high transduction efficiency, we constructed recombinant PHP.eB adeno-associated virus (AAV) vectors expressing VP16 proteins of PRV or HSV. Since we did not want to affect protein-protein interactions, we designed a cleavable fluorophore-VP16 construct (mTurqoise-P2A-VP16) instead of a fusion protein. Upon transduction of neurons with these AAV vectors, abundant amounts of full-length and cleaved proteins were detected after 4 days. When the axons of these neurons were infected with a latency establishing dose of PRV180, we found that HSV-1 VP16 expression was sufficient to induce a fast escape from genome silencing. However, PRV VP16 was not able to induce productive infection. Moreover, HSV VP16 expression successfully reactivated latent PRV infection while expression of PRV VP16 did not. These results were surprising not only because PRV VP16 expression was not able to activate PRV productive infection but also, because HSV-1 VP16 is known to have a weak affinity to murine oct-1 [42]. Therefore, we did not expect a strong activation of PRV IE gene expression leading to fast escape from silencing.

To explore the indirect activity of HSV-1 VP16 on PRV gene expression we identified the genes in neurons whose transcription was increased after transduction with HSV or PRV VP16 expressing AAVs without herpesvirus infection. We analyzed differential gene expression profiles of HSV-VP16 and PRV-VP16 transduced neurons in comparison to the mTurqoise2 expressing AAV transduced ones by RNA seq experiment. We found that the gene expression profiles were surprisingly different in HSV-VP16 versus PRV VP16 expressing neurons.

Transcription of the Dusp4 and Adcyap1 genes was increased significantly in PRV VP16 transduced neurons. Dual specificity phosphatase 4 (Dusp4) or Mitogen activated protein kinase phosphatase 2 (MKP-2), is a dual specific nuclear phosphatase that is selective for both ERK and JNK. These phosphatases regulate the activities of the MAP kinases through dual specific dephosphorylation at tyrosine and threonine residues fine tuning the amplitude and extent of cellular activation. Cadalbert et al. showed that Dusp4 has specificity for JNK (not ERK) in vivo [47]. We did not detect JNK activation in neurons when PRV VP16 was expressed (as we detected with HSV VP16 transduction) and the elevated levels of Dusp4 might be the reason for that. Interestingly, the gene with the second highest expression level was Adcyap1 (Adenylate cyclase activating polypeptide 1, i.e., PACAP) gene. This gene encodes a secreted pro-protein that is further processed into multiple peptides that stimulate adenylate cyclase and increase cyclic adenosine monophosphate (cAMP) levels. We have previously shown that increased cAMP levels in neurons either upon forskolin or cell permeable dibutryl cAMP treatment induces slow PRV escape from gene silencing [8]. Surprisingly, PRV VP16 protein expression in neurons, even with increased Adcyap1 transcription, was not sufficient to induce PRV escape from silencing. Our data suggests that PRV VP16, unlike the HSV-1 homolog, requires an additional viral co-factor to interact with neuronal factors and activate viral transcription to enable PRV escape from genome silencing.

A clear signature of HSV VP16 expression in primary neurons is the significant activation of proto-oncogene expression dominated by Jun, JunD and Pim2 genes. Jun proteins can induce cell transformation dependent on phosphorylation by the c-Jun N-terminal kinase (JNK). We have previously reported that the activity of JNK is essential in neuronal stress mediated PRV escape from genome silencing [8]. Active JNK was not essential in viral tegument mediated escape from genome silencing in the same model. Therefore, we hypothesized that HSV VP16 mediated escape from PRV genome silencing is mediated by elevated Jun activity. In support of this hypothesis, we detected increased levels of Jun and phosphorylated Jun protein only when neurons were transduced with HSV-16 expressing AAVs. Furthermore, when we inhibited Jun phosphorylation using various JNK inhibitors, we consistently decreased the efficiency of HSV VP16 mediated PRV escape from genome silencing. Cliffe et al. also demonstrated that JNK activity is critical for reactivation of HSV-1; particularly for lytic gene expression during the initial phase (phase I) of reactivation [22, 28]. These results confirm that HSV VP16 activity depends on active JNK similar to the cellular stress pathway mediated but differs from PRV tegument mediated fast escape from genome silencing. However, the HSV-VP16 mediated escape happens rapidly in the neuronal cell bodies with kinetics resembling the tegument mediated fast escape. This might require the action of other proteins with elevated expression levels.

c-Jun, together with its interaction partner c-Fos, forms the AP-1 transcription factor complex. Increased levels of c-Jun and c-Fos, and increased activity of the AP-1 complex was observed in response to neuronal activity [48]. Interestingly, the AP-1 complex was shown to be necessary to activate the major immediate early promoter (MIEP) of human cytomegalovirus (HCMV), and a viral GPCR (pUs28) was identified for interfering with Fos activation to shift the infection mode toward latency in human leukemia monocytic cells [49]. Although we detected increased levels of Jun and phosphorylated Jun in HSV VP16 transduced neurons, we did not detect significant difference in Fos transcripts.

The Pim2 gene had the second highest enrichment in HSV VP16 transduced neurons. The proviral integration site for Moloney murine leukemia virus 2 (Pim2) is a serine/threonine kinase belonging to the PIM family of kinases and plays an important role as an oncogene in multiple cancers including leukemia, myeloma, prostate, and breast cancers. PIM2 is largely expressed in both leukemia and solid tumors, and it promotes the transcriptional activation of genes involved in cell survival, cell proliferation, and cell-cycle progression. The effect of Pim2 on VP16 mediated activation of alpha herpesvirus gene transcription whether it is necessary as Jun activation requires further investigation.

Although the VP16 proteins of HSV-1 and PRV induced different gene expression profiles in neurons, the transcription of two genes was reduced for both. These genes are argininosuccinate synthetase 1 (Ass1) and solute carrier family 18 member A3 (Slc18a3). Ass1, as a homotetramer, catalyzes the synthesis of argininosuccinate from aspartate and citrulline. This enzyme primarily functions as the limiting step in the citrulline-nitric oxide (NO) cycle in many cell types. Grady et al., found that Ass1 knockdown by siRNA increases HSV-1 yield in fibroblasts [50]. They also showed that Ass1-deficient cells make more viral genomes, express higher levels of viral mRNA and proteins, and exhibit the metabolic signature of HSV-1 infected fibroblasts. It is interesting that we observed reduced transcription of this gene in primary neurons upon expression of both homologs VP16 proteins. As evidenced by Grady et al., such metabolic state is preferable to support viral DNA replication, mRNA, and protein expression during a productive infection. It appears that VP16 protein, as a component of the incoming virions, not only interacts with host cell transcription factors to initiate viral gene expression, but also primes host cell metabolism for productive infection. Vesicular acetylcholine transporter (SLC18A3) mediates the uptake of acetylcholine (Ach), a classical neurotransmitter. Elevated ACh was shown to activate the protein kinase A (PKA)/cAMP-response element binding protein (CREB) pathway, a key pathway in stress mediated escape from genome silencing and reactivation of alpha herpesviruses. Surprisingly we detected reduced expression of this transporter after expression of both HSV and PRV VP16 in neurons. It would be interesting to test the effect of the knock-down of Slc18a3 on yield of PRV and HSV-1 in neurons.

In summary, our results showed that prior HSV-1 VP16 expression in neurons enabled incoming PRV genomes to escape from silencing primarily by activating Jun signaling. However, this escape is not a slow escape as we observed with cellular stress mediated pathway that depends on active PKA and JNK kinases. HSV VP16 expressing neurons start accumulating the PRV capsid protein in 2 days, and infection spreads throughout the whole S-compartments in 3 days. Such fast escape was observed when PRV tegument proteins were present in the S-compartments that alleviated the need for active cellular kinases such as PKA and JNK. Surprisingly, we did not detect escape from silencing or reactivation events upon PRV VP16 expression in neurons leading us to hypothesize that PRV VP16 requires a viral co-factor to initiate IE180 transcription. Our future studies will aim to identify the interaction partners of HSV and PRV VP16 in primary neurons with or without virus infection.

## Materials and Methods

### Cells Lines and Viruses

The cell lines used in this study are from ATCC; Rat2 (rat fibroblasts), PK15 (pig kidney epithelial cells) and Vero cells (African green monkey kidney cells). All cell lines were grown in DMEM media containing 10% fetal bovine serum (FBS). The virus strains used in this study were HSV-1 OK14 and PRV 180. HSV-1 OK14 was constructed by co-transfecting BamHI-digested pHSV1(17+) Lox-mRFPVP26 and purified HSV-1(17+) DNA (O. Kobiler and L. W. Enquist, unpublished data). The pHSV1(17+) Lox-mRFPVP26 was a kind gift from Katinka Döhner and Beate Sodeik. PRV 180 expresses an mRFP-VP26 fusion protein in a PRV-Becker background [51].

### Primary Neuron Cultures

Experiments on neurons were performed using primary superior ganglionic (SCG) neuron cultures. The following cell culture procedure is adapted from Ch’ng and Enquist, and lasts between 14-21 days in total [52]. SCG neurons isolated from Sprague-Dawley rat embryos (Hilltop Labs) were cultured in either 35-mm, 12-well, or 6-well dishes. After coating the culture dishes with poly-DL-ornithine (Sigma) and natural mouse laminin (Invitrogen), 2/3 of tryspsinized and triturated ganglion was plated per S-compartment. The neuronal medium used was supplemented with nerve growth factor 2.5S (Invitrogen), B27 (Gibco), penicillin and streptomycin. After 2 to 3 days, cytosine-D-arabinofuranoside (Ara-C; Sigma) was added for a minimum of 2 days to kill non-neuronal cells. All animal work was done in accordance with the Institutional Animal Care and Use Committee of the Princeton University Research Board under protocol 1947-13

### Recombinant Adeno Associated Virus (AAV) Vectors

The AAV vectors are designed to express HSV-1 VP16 (AAV_mTurq_HSV VP16) or PRV VP16 (AAV_mTurq_PRV VP16) (Figure 1). They encode a polyprotein containing a self-cleaving picornavirus P2A cleavage site in the middle. One part of the polyprotein consists of the VP16 protein, either from HSV-1 or PRV, and on the other side of the P2A site, the AAV encodes a diffusible fluorophore: mTurqoise2. When the P2A site cleaves, it cleaves at its C-terminus, leading only one amino acid attached to the VP16 protein. This fluorophore serves as a reporter to test the transduction efficiency of the AAV vector. To reduce the risk of the reporter interfering with the function of VP16, a cleavable polyprotein was chosen over the use of a fusion protein. The SCGs that were transduced with these AAV vectors were plated in 6-well dishes and the following amounts of AAV vector were added to each well: 8×10^11^ genome copies of AAV_mTurq, 1.9×10^12^ genome copies of AAV_HSV_VP16, and 1.4×10^12^ genome copies of AAV_PRV_VP16 for both RNA-seq and IF analysis. All PHP.eB AAVs were constructed by the Princeton Neuroscience Institute Viral Core Facility using triple plasmid transfection of HEK 293 cells as previously described [53]. The pUCmini-iCAP-PHP.eB was a gift from Viviana Gradinaru (Addgene plasmid # 103005; http://n2t.net/addgene:103005 ; RRID:Addgene_103005).

### Antibodies and Inhibitors

The main primary antibodies used for gel analysis of virus-infected cells were a commercial HSV-1 VP16 antibody (Abcam) and a rabbit polyclonal antibody against the full-length PRV VP16 that was ordered from Genscript. Anti-cJun and anti-phospho-cJun (Ser63) antibodies were purchased from Cell Signaling. Goat-anti-mouse and goat-anti-rabbit, horseradish peroxidase (HRP) secondary antibodies were used for their appropriate primary antibodies at 1:5000 dilution (KPL). Fluorescent anti-rabbit and anti-mouse Alexa-488 and Alexa-546 secondary antibodies from Invitrogen were used for IF. SP600125 was ordered from Sigma (420119) (used at 10 μM), JNK-IN-8 was ordered from Sellechem (ab145194) (used at 5 μM), AS601245 was ordered from Abcam (ab145194) (used at 20 μM).

### Immunofluorescence (IF) analysis

To prepare cells for IF, they were grown either on glass coverslips in 6-well dishes or in 8-well chamber slides. Cells were fixed using 4% paraformaldehyde (PFA) at room temperature or methanol at -20°C. Blocking was done using a 3% milk powder solution in PBS-Tween (0.05%) for one hour. Primary and secondary antibody dilutions were both prepared in 1% milk powder solution in PBS-Tween. The primary antibody staining was done for one hour at room temperature without rocking and then washed three times with PBS-Tween while rocking. Secondary antibody (Alexa fluor) staining was also done for one hour at room temperature with the dishes covered. After three more washes in PBS-Tween rocking at room temperature, the cells grown on coverslips were mounted onto glass microscope slides. Images were taken of AAV-transduced or PRV-infected SCGs and the IF patterns of VP16-directed fluorescent antibodies compared. Images were taken using a Nikon Eclipse Ti fluorescence microscope and analyzed using NIS Elements software.

### Polyacrylamide Gel Electrophoresis and Western Blot Analysis

After infection with virus, Vero and Rat2 cells were collected after set amounts of time and protein lysates extracted. These protein lysates were analyzed by SDS polyacrylamide gel electrophoresis (PAGE) and Western blotting (WB). To run SDS-PAGE, proteins were extracted from the supernatant of lysed cells. This Western blot (WB) procedure is adapted from [9]. Cells were lysed in a mixture of radioimmunoprecipitation assay (RIPA-light) buffer, 1 mM dithiothreitol (DTT), and protease inhibitor cocktail (Sigma-Aldrich). After incubation on ice for 30 minutes, followed by sonication, and centrifugation, the supernatants were extracted, mixed with 5x Laemmli buffer, and heated at 90°C for 5 minutes before running the gel. SDS-PAGE was performed using 4-12% polyacrylamide gels. After running the gel, the proteins were transferred from the gel onto nitrocellulose membranes. Blocking was performed with 5% milk powder in PBS for 1 hour at room temperature while rocking or overnight at 4ºC while rocking. The blots were transferred to nitrocellulose membranes and immunoblots performed with primary and secondary antibodies. Membranes were incubated first with primary antibodies for 1 hour at room temperature or overnight at 4°C and then with HRP-coupled secondary antibodies (KPL) for 1 hour at room temperature. After washing, the protein bands were visualized by exposure on HyBlot CL (Denville scientific) blue x-ray films. For the Vero infection and Rat2 infection time-course, cells were grown in 6-well dishes with two wells being harvested together and loaded for each lane of SDS-PAGE.

### RNA-Sequencing and data analysis

RNA-seq was used to compare the cellular gene activation levels of SCG neurons transduced with PRV VP16, HSV-1 VP16, and mTurqoise2 vectors. The SCGs cultured in 6-well dishes and one entire 6-well was harvested and pooled together for RNA isolation (RNeasy, Qiagen) per sample. This was repeated three times per sample and RNA-seq libraries were prepared in triplicates. The following library preparation and sequencing procedures are adapted from Kovanda et al. and were performed with the help of the microarray facility [54]. To prepare samples for RNA-seq, multiple samples of RNA were first isolated from AAV-transduced SCGs. The RNA concentration and RNA integrity number (RIN) of each sample were measured to determine which samples are of high enough quality to build the small RNA library needed for sequencing. Once the small RNA libraries were produced using a prep kit, the corresponding single-stranded complementary DNAs (cDNAs) were PCR amplified and the PCR products purified and size-selected with gel purification. Purified PCR products within a given size range were sequenced (75 nucleotide single end sequencing). RNA sequencing (RNA-seq) was performed on superior cervical ganglia (SCG) neurons transduced with one of three AAV vectors: AAV_mTurqoise2 (mTurqoise2 alone), AAV_PRV_VP16 (PRV UL48 polyprotein), and AAV_HSV_VP16 (HSV VP16 polyprotein). RNA-seq was run by the High-Throughput Sequencing and Microarray facility (Princeton University) and analysis of RNA-Seq data was performed with the help of Lance Parsons from Princeton University Lewis Sigler Institute. Analysis consisted of four main steps: i) Alignment of the raw reads to the reference genome using the STAR aligner ii) A count of the reads aligned to each gene using featureCounts iii) QC on the raw data, alignment, and quantification using FastQC, samtools, RSeQC, and MultiQC iv) Differential expression analysis using DESeq2. The reference genome used was a combination of the *Rattus norvegicus* genome with additional sequences for mTurqoise2, HSV VP16 and PRV UL48. After running differential expression (DESeq2) analysis, RUVseq (Remove Unwanted Variation) was used to remove variation due to batch effects. To further analyze differential gene expression, the log2-fold change (logFC) and the adjusted P-value (p-adj) of each gene in the DESeq2 results were prioritized. A p-adj value of 0.5 was set as the cutoff and any gene with p-adj > 0.5 was disregarded. The p-adj value was calculated by taking the p-value derived from a Wald test and using the Benjamini-Hochberg procedure to control for false discovery rate (FDR).

<u>Supplemental excel sheet.</u> HSV vs. PRV VP16 differential gene expression

## References

1. Looker, K.J., et al., The global and regional burden of genital ulcer disease due to herpes simplex virus: a natural history modelling study. BMJ Glob Health, 2020. 5(3): p. e001875.

2. Looker, K.J., et al., Global and Regional Estimates of Prevalent and Incident Herpes Simplex Virus Type 1 Infections in 2012. PLoS One, 2015. 10(10): p. e0140765.

3. McCormick, I., et al., Incidence of Herpes Simplex Virus Keratitis and Other Ocular Disease: Global Review and Estimates. Ophthalmic Epidemiol, 2022. 29(4): p. 353–362.

4. Koyuncu, O.O., I.B. Hogue, and L.W. Enquist, Virus infections in the nervous system. Cell Host Microbe, 2013. 13(4): p. 379–93.

5. Looker, K.J., et al., First estimates of the global and regional incidence of neonatal herpes infection. Lancet Glob Health, 2017. 5(3): p. e300–e309.

6. Antinone, S.E. and G.A. Smith, Retrograde axon transport of herpes simplex virus and pseudorabies virus: a live-cell comparative analysis. J Virol, 2010. 84(3): p. 1504–12.

7. Hafezi, W., et al., Entry of herpes simplex virus type 1 (HSV-1) into the distal axons of trigeminal neurons favors the onset of nonproductive, silent infection. PLoS Pathog, 2012. 8(5): p. e1002679.

8. Koyuncu, O.O., et al., Compartmented neuronal cultures reveal two distinct mechanisms for alpha herpesvirus escape from genome silencing. PLoS Pathog, 2017. 13(10): p. e1006608.

9. Koyuncu, O.O., et al., The number of alphaherpesvirus particles infecting axons and the axonal protein repertoire determines the outcome of neuronal infection. mBio, 2015. 6(2).

10. Fan, D., et al., The Role of VP16 in the Life Cycle of Alphaherpesviruses. Front Microbiol, 2020. 11: p. 1910.

11. Sawtell, N.M. and R.L. Thompson, De Novo Herpes Simplex Virus VP16 Expression Gates a Dynamic Programmatic Transition and Sets the Latent/Lytic Balance during Acute Infection in Trigeminal Ganglia. PLoS Pathog, 2016. 12(9): p. e1005877.

12. Wysocka, J. and W. Herr, The herpes simplex virus VP16-induced complex: the makings of a regulatory switch. Trends Biochem Sci, 2003. 28(6): p. 294–304.

13. Campbell, M.E., J.W. Palfreyman, and C.M. Preston, Identification of herpes simplex virus DNA sequences which encode a trans-acting polypeptide responsible for stimulation of immediate early transcription. J Mol Biol, 1984. 180(1): p. 1–19.

14. Wilson, A.C., et al., Combinatorial control of transcription: the herpes simplex virus VP16-induced complex. Cold Spring Harb Symp Quant Biol, 1993. 58: p. 167–78.

15. Antinone, S.E., et al., The Herpesvirus capsid surface protein, VP26, and the majority of the tegument proteins are dispensable for capsid transport toward the nucleus. J Virol, 2006. 80(11): p. 5494–8.

16. Bohannon, K.P., et al., Differential protein partitioning within the herpesvirus tegument and envelope underlies a complex and variable virion architecture. Proc Natl Acad Sci U S A, 2013. 110(17): p. E1613–20.

17. Buch, A., et al., Inner tegument proteins of Herpes Simplex Virus are sufficient for intracellular capsid motility in neurons but not for axonal targeting. PLoS Pathog, 2017. 13(12): p. e1006813.

18. Radtke, K., et al., Plus- and minus-end directed microtubule motors bind simultaneously to herpes simplex virus capsids using different inner tegument structures. PLoS Pathog, 2010. 6(7): p. e1000991.

19. Wolfstein, A., et al., The inner tegument promotes herpes simplex virus capsid motility along microtubules in vitro. Traffic, 2006. 7(2): p. 227–37.

20. Roizman, B. and A.E. Sears, An inquiry into the mechanisms of herpes simplex virus latency. Annu Rev Microbiol, 1987. 41: p. 543–71.

21. Thompson, R.L. and N.M. Sawtell, Targeted Promoter Replacement Reveals That Herpes Simplex Virus Type-1 and 2 Specific VP16 Promoters Direct Distinct Rates of Entry Into the Lytic Program in Sensory Neurons in vivo. Front Microbiol, 2019. 10: p. 1624.

22. Cliffe, A.R. and A.C. Wilson, Restarting Lytic Gene Transcription at the Onset of Herpes Simplex Virus Reactivation. J Virol, 2017. 91(2).

23. Wilson, A.C. and I. Mohr, A cultured affair: HSV latency and reactivation in neurons. Trends Microbiol, 2012. 20(12): p. 604–11.

24. Muller, T., et al., Pseudorabies virus in wild swine: a global perspective. Arch Virol, 2011. 156(10): p. 1691–705.

25. Heine, J.W., et al., Proteins specified by herpes simplex virus. XII. The virion polypeptides of type 1 strains. J Virol, 1974. 14(3): p. 640–51.

26. Chan, K.Y., et al., Engineered AAVs for efficient noninvasive gene delivery to the central and peripheral nervous systems. Nat Neurosci, 2017. 20(8): p. 1172–1179.

27. Maturana, C.J., et al., Novel tool to quantify with single-cell resolution the number of incoming AAV genomes co-expressed in the mouse nervous system. Gene Ther, 2021.

28. Cliffe, A.R., et al., Neuronal Stress Pathway Mediating a Histone Methyl/Phospho Switch Is Required for Herpes Simplex Virus Reactivation. Cell Host Microbe, 2015. 18(6): p. 649–58.

29. Cuddy, S.R., et al., Neuronal hyperexcitability is a DLK-dependent trigger of herpes simplex virus reactivation that can be induced by IL-1. Elife, 2020. 9.

30. Roizman, B. and R.J. Whitley, An inquiry into the molecular basis of HSV latency and reactivation. Annu Rev Microbiol, 2013. 67: p. 355–74.

31. Cliffe, A.R. and A.C. Wilson, Restarting Lytic Gene Transcription at the Onset of Herpes Simplex Virus Reactivation. J Virol, 2016.

32. Koyuncu, O.O., M.A. MacGibeny, and L.W. Enquist, Latent versus productive infection: the alpha herpesvirus switch. Future Virol, 2018. 13(6): p. 431–443.

33. Luxton, G.W., et al., Targeting of herpesvirus capsid transport in axons is coupled to association with specific sets of tegument proteins. Proc Natl Acad Sci U S A, 2005. 102(16): p. 5832–7.

34. Suk, H. and D.M. Knipe, Proteomic analysis of the herpes simplex virus 1 virion protein 16 transactivator protein in infected cells. Proteomics, 2015. 15(12): p. 1957–67.

35. Goodrich, J.A., et al., Drosophila TAFII40 interacts with both a VP16 activation domain and the basal transcription factor TFIIB. Cell, 1993. 75(3): p. 519–30.

36. Klemm, R.D., et al., Molecular cloning and expression of the 32-kDa subunit of human TFIID reveals interactions with VP16 and TFIIB that mediate transcriptional activation. Proc Natl Acad Sci U S A, 1995. 92(13): p. 5788–92.

37. Langlois, C., et al., NMR structure of the complex between the Tfb1 subunit of TFIIH and the activation domain of VP16: structural similarities between VP16 and p53. J Am Chem Soc, 2008. 130(32): p. 10596–604.

38. Mittler, G., et al., A novel docking site on Mediator is critical for activation by VP16 in mammalian cells. EMBO J, 2003. 22(24): p. 6494–504.

39. Yang, F., et al., The activator-recruited cofactor/Mediator coactivator subunit ARC92 is a functionally important target of the VP16 transcriptional activator. Proc Natl Acad Sci U S A, 2004. 101(8): p. 2339–44.

40. Thompson, R.L., C.M. Preston, and N.M. Sawtell, De novo synthesis of VP16 coordinates the exit from HSV latency in vivo. PLoS Pathog, 2009. 5(3): p. e1000352.

41. Diefenbach, R.J., et al., Transport and egress of herpes simplex virus in neurons. Rev Med Virol, 2008. 18(1): p. 35–51.

42. Kim, J.Y., et al., Transient reversal of episome silencing precedes VP16-dependent transcription during reactivation of latent HSV-1 in neurons. PLoS Pathog, 2012. 8(2): p. e1002540.

43. Miranda-Saksena, M., et al., Anterograde transport of herpes simplex virus type 1 in cultured, dissociated human and rat dorsal root ganglion neurons. J Virol, 2000. 74(4): p. 1827–39.

44. Roizman, B., G. Zhou, and T. Du, Checkpoints in productive and latent infections with herpes simplex virus 1: conceptualization of the issues. J Neurovirol, 2011. 17(6): p. 512–7.

45. Fuchs, W., et al., The UL48 tegument protein of pseudorabies virus is critical for intracytoplasmic assembly of infectious virions. J Virol, 2002. 76(13): p. 6729–42.

46. Fuchs, W., H. Granzow, and T.C. Mettenleiter, A pseudorabies virus recombinant simultaneously lacking the major tegument proteins encoded by the UL46, UL47, UL48, and UL49 genes is viable in cultured cells. J Virol, 2003. 77(23): p. 12891–900.

47. Cadalbert, L., et al., Conditional expression of MAP kinase phosphatase-2 protects against genotoxic stress-induced apoptosis by binding and selective dephosphorylation of nuclear activated c-jun N-terminal kinase. Cell Signal, 2005. 17(10): p. 1254–64.

48. Yap, E.L. and M.E. Greenberg, Activity-Regulated Transcription: Bridging the Gap between Neural Activity and Behavior. Neuron, 2018. 100(2): p. 330–348.

49. Krishna, B.A., et al., Human cytomegalovirus G protein-coupled receptor US28 promotes latency by attenuating c-fos. Proc Natl Acad Sci U S A, 2019. 116(5): p. 1755–1764.

50. Grady, S.L., et al., Argininosuccinate synthetase 1 depletion produces a metabolic state conducive to herpes simplex virus 1 infection. Proc Natl Acad Sci U S A, 2013. 110(51): p. E5006–15.

51. del Rio, T., et al., Heterogeneity of a fluorescent tegument component in single pseudorabies virus virions and enveloped axonal assemblies. J Virol, 2005. 79(7): p. 3903–19.

52. Ch’ng, T.H. and L.W. Enquist, Neuron-to-cell spread of pseudorabies virus in a compartmented neuronal culture system. J Virol, 2005. 79(17): p. 10875–89.

53. Chan, A., C.J. Maturana, and E.A. Engel, Optimized formulation buffer preserves adeno-associated virus-9 infectivity after 4 degrees C storage and freeze/thawing cycling. J Virol Methods, 2022. 309: p. 114598.

54. Kovanda, A., et al., Differential expression of microRNAs and other small RNAs in muscle tissue of patients with ALS and healthy age-matched controls. Sci Rep, 2018. 8(1): p. 5609.

